# DNA Barcoding of fogged caterpillars in Peru: A novel approach for unveiling host-plant relationships of tropical moths (Insecta, Lepidoptera)

**DOI:** 10.1101/799221

**Authors:** Axel Hausmann, Juliane Diller, Jerome Moriniere, Amelie Höcherl, Andreas Floren, Gerhard Haszprunar

## Abstract

A total of 130 lepidopteran larvae were selected from 37 fogging samples at the Panguana station, district Yuyapichis, province Puerto Inca, department Huánuco, Peru. Target trees were pre-identified and subsequently submitted to molecular confirmation of identity with three markers (rbcL, psbA and trnL-F). Identification of 119 lepidopteran larvae (92 species) was successful through DNA barcoding: Comparison of COI barcodes with the reference database of adult moths resulted in 65 (55%) matches at species level, 32 (27%) at genus level and 19 (16%) at subfamily or family level. Three larvae could not be assigned to a family. For these larvae the fogged target tree now suggests a potential host-plant relationship. Molecular gut content analysis, based on High-Throughput-Sequencing was successfully tested for ten larvae corroborating feeding on the target plant in some cases but elucidating several other cases of potential ‘alternative feeding’. We propose a larger-scale approach using this rapid and efficient method including molecular gut-content analyses for comprehensively testing the ratio of ‘alternative feeders’ and pitfalls caused by collateral fogging of larvae from neighboring trees.

## Introduction

Despite much valuable work on host-relationships of Neotropical moths, e.g. from Ecuador [1], [2], or Costa Rica [3], [4], [5], the relevant literature is still scarce and patchy compared with the huge species diversity of Lepidoptera in Central and South America. Apart from the aforementioned programs only few original data are published for host-plant relationships of Lepidoptera and much of the work focused on caterpillars found on plants of economic importance (pests and potential pests) (e.g. [6], [7], [8]).

Exemplified from one of the most diverse moth families, Geometridae, the largest project in Costa Rica so far revealed the huge amount of 22,957 geometrid moth records, the barcoded reared adults clustering to 566 BINs, of which 162 currently having Linnean species names (D.J. Janzen & W. Hallwachs pers. comm.). Brehm [1] presented 48 neotropical geometrid species with host-plant records, with 11 records added by Dyer et al. [9] and 59 records by Bodner et al. [2]. Thus, altogether for some 680 Neotropical geometrid species (about 270 of which with Linnean species names) host-plant relationships are known, covering approx. 8-10% (4% with Linnean names) of the described geometrid fauna of Central and South America (for estimations of total number of described geometrid species cf. Scoble et al. [10]: 6433 species; Heppner [11]: 7956 species).

Estimation for Neotropical species diversity is based on approx. 37,000 described Neotropical moth species (Heppner [11]: 44,800 described Lepidoptera, including approx. 7800 Rhopalocera species [12]) and considering that (a) ‘Microlepidoptera’ are severely understudied and (b) the vast majority of the Neotropical moth fauna is still undescribed as suggested by the ratio of undescribed species in some 380,000 Neotropical lepidopteran DNA barcodes on Barcode of Life Data Systems (‘BOLD’). Extrapolating the aforementioned data on species numbers and feeding records we estimate that for >98% of the putatively >100,000 Neotropical moth species authentic feeding records from nature are lacking.

Traditionally, most insect larvae are identified by rearing them to the adult stage and by analysing the morphology of the adult. Methodological constraints in this classic approach are (1) visual search and collecting on plant depending on the skills of the biologist, (2) the canopy region of trees hardly accessible, (3) nocturnal activity of many larvae requiring difficult search by night, (4) collecting without feeding observation may lead to misinterpretations ([13], [14]: 20-50%, “alternative feeders” on lichens, dead leaves, algae, etc), (5) beating, shaking, net-sweeping may obscure the real where-about of the larva, (6) feeding records in rearing may not reflect the natural host-plant association, (7) rearing to adult is time consuming, (8) rearing may fail (deseases, parasitoids), (9) identification and availability of host-plant (for rearing) often difficult.

Molecular identification of lepidopteran larvae and other insects through DNA barcoding (COI 5’) was repeatedly carried out successfully [15], [16], [17], [18], [19], permitting an easy, cheap and rapid identification of larvae collected from their host-plants. Identification through DNA barcoding is possible even from dry skins after moulting and from empty pupal exuviae after hatching of the moths (own, unpublished data). Currently, there are large-scale projects devoted to the identification of larvae along with their host-plants in Papua New Guinea ([20]) and Costa Rica ([4]). Both are based on an integrative approach combining morphology, rearing and molecular techniques for the identification of the reared adults and/or their parasitoids.

Miller et al. [15] and Matheson et al. [16] investigated and ascertained relationships between plants and caterpillars through a method based on the DNA identification of the larval gut content, an effective but (in earlier times) expensive and time-consuming approach, especially as a routine application in larger surveys. Later on, molecular gut content analysis was proposed for unveiling insect-host plant associations e.g. for beetles [21], [22], [23], [24], and for soil insects [25].

The aim of this pilot paper was to establish methodology to infer host-plant relationships of caterpillars based on the identification of larvae collected by insecticidal knock-down (canopy-fogging) on their food-plants through DNA barcoding and to use gut HTS-based content analysis to estimate potential pitfalls due to ‘alternative feeding’ or due to collateral fogging from neighbouring plants, lianas etc. (cf. discussion).

## Material and Methods

### Collecting/Canopy fogging

Canopy fogging was performed by AmH und AF from the ground with a Swingfog SN 50 fogger, using natural Pyrethrum, diluted in a highly raffinated white oil, as knock-down agent to prevent the introduction of persistent chemicals into the environment. For details of the fogging procedure see [26]. In most cases, trees with dense foliage cover and little canopy overlap with neighboring trees were chosen. We made sure the fog reached the canopy and stood there for at least five minutes to affect the arthropods. In order to install the collecting sheets, larger saplings and other interferring vegetation elements were cleared below the tree projection area. All organisms dropping down from the trees were collected at least one hour after the fogging from expanded plastic sheets of 20m^2^ size, covering an estimated minimum of 80% of the target tree canopy. The arthropods were then transferred into accurately labelled jars with 100% ethanol without pre-sorting. The following day the Ethanol was renewed and excessive plant material with its high water content was removed. Samples were stored at room temperature for up to two weeks while in the research station Panguana. The ethanol was renewed again when the samples were added to the Zoologische Staatssammlung München in Bavaria, Germany (SNSB – ZSM).

The study site is located in the westernmost Amazonian Basin, eastern central Peru, department Huánuco, at the ACP Panguana station (−9.613393°N −74.935911°E; 222 m; see also [27]), the fogged target trees were all situated in a radius of less than 2000 metres around the station. Collecting was performed in the late afternoon, betwen 17 and 19 o clock, from 24^th^ of November to 8^th^ of December 2017. For identification of the target trees see results.

### Tissue sampling and identification of larvae (DNA barcoding, COI)

Out of 47 samples – each of them referring to a fogged tree (cf. Table 3 and Supporting information S1 Table) – all lepidopteran larvae were separated, in total 130 specimens. The larvae were dried on paper, photographed and then separately stored in Eppendorf tubes. A list of all 130 larvae along with their fogging sample number is given in Table 1, examples are shown in Figure 1. Tissue sampling was carried out for all 130 larvae by using scissors and pincers, which were carefully cleaned after each tissue sampling in 100% alcohol to avoid contamination among samples. Tissues (one vertically cut segment, in very small larvae two segments) were transferred to a lysis plate under 0.5 ml 100% alcohol.

**Table 1.** Overview on the 130 Lepidoptera larvae selected from 36 fogging samples from Panguana, Peru, sequencing success and number of target tree.

| Larva Nr. | Barcode ID | Sequence Length (bp) | Target tree nr. |
| --- | --- | --- | --- |
| 1 | BC ZSM Lep 98047 | 658 | 2 |
| 2 | BC ZSM Lep 98048 | 658 | 2 |
| 3 | BC ZSM Lep 98049 | 658 | 2 |
| 4 | BC ZSM Lep 98050 | 658 | 3 |
| 5 | BC ZSM Lep 98051 | 658 | 7 |
| 6 | BC ZSM Lep 98052 | 658 | 9 |
| 7 | BC ZSM Lep 98053 | 658 | 9 |
| 8 | BC ZSM Lep 98054 | 658 | 14 |
| 9 | BC ZSM Lep 98055 | 658 | 14 |
| 10 | BC ZSM Lep 98056 | 658 | 15 |
| 11 | BC ZSM Lep 98057 | 658 | 16 |
| 12 | BC ZSM Lep 98058 | 658 | 16 |
| 13 | BC ZSM Lep 98059 | 658 | 19 |
| 14 | BC ZSM Lep 98060 | 658 | 20 |
| 15 | BC ZSM Lep 98061 | 635 | 20 |
| 16 | BC ZSM Lep 98062 | 658 | 20 |
| 17 | BC ZSM Lep 98063 | 658 | 21 |
| 18 | BC ZSM Lep 98064 | 658 | 22 |
| 19 | BC ZSM Lep 98065 | 658 | 22 |
| 20 | BC ZSM Lep 98066 | 658 | 23 |
| 21 | BC ZSM Lep 98067 | 658 | 23 |
| 22 | BC ZSM Lep 98068 | 658 | 24 |
| 23 | BC ZSM Lep 98069 | 658 | 24 |
| 24 | BC ZSM Lep 98070 | 658 | 25 |
| 25 | BC ZSM Lep 98071 | 0 | 27 |
| 26 | BC ZSM Lep 98072 | 658 | 27 |
| 27 | BC ZSM Lep 98073 | 658 | 27 |
| 28 | BC ZSM Lep 98074 | 658 | 27 |
| 29 | BC ZSM Lep 98075 | 658 | 28 |
| 30 | BC ZSM Lep 98076 | 658 | 28 |
| 31 | BC ZSM Lep 98077 | 658 | 29 |
| 32 | BC ZSM Lep 98078 | 658 | 32 |
| 33 | BC ZSM Lep 98079 | 658 | 32 |
| 34 | BC ZSM Lep 98080 | 658 | 32 |
| 35 | BC ZSM Lep 98081 | 658 | 32 |
| 36 | BC ZSM Lep 98082 | 658 | 32 |
| 37 | BC ZSM Lep 101897 | 658 | 1 |
| 38 | BC ZSM Lep 101898 | 658 | 1 |
| 39 | BC ZSM Lep 101899 | 658 | 1 |
| 40 | BC ZSM Lep 101900 | 658 | 2 |
| 41 | BC ZSM Lep 101901 | 0 | 2 |
| 42 | BC ZSM Lep 101902 | 658 | 2 |
| 43 | BC ZSM Lep 101903 | 658 | 5 |
| 44 | BC ZSM Lep 101904 | 0 | 7 |
| 45 | BC ZSM Lep 101905 | 658 | 7 |
| 46 | BC ZSM Lep 101906 | 658 | 9 |
| 47 | BC ZSM Lep 101907 | 658 | 9 |
| 48 | BC ZSM Lep 101908 | 658 | 10 |
| 49 | BC ZSM Lep 101909 | 658 | 10 |
| 50 | BC ZSM Lep 101910 | 658 | 10 |
| 51 | BC ZSM Lep 101911 | 658 | 10 |
| 52 | BC ZSM Lep 101912 | 658 | 11 |
| 53 | BC ZSM Lep 101913 | 658 | 12 |
| 54 | BC ZSM Lep 101914 | 658 | 12 |
| 55 | BC ZSM Lep 101916 | 658 | 13 |
| 56 | BC ZSM Lep 101917 | 658 | 13 |
| 57 | BC ZSM Lep 101918 | 0 | 13 |
| 58 | BC ZSM Lep 101919 | 658 | 14 |
| 59 | BC ZSM Lep 101920 | 658 | 14 |
| 60 | BC ZSM Lep 101921 | 658 | 14 |
| 61 | BC ZSM Lep 101922 | 658 | 14 |
| 62 | BC ZSM Lep 101923 | 658 | 14 |
| 63 | BC ZSM Lep 101924 | 658 | 14 |
| 64 | BC ZSM Lep 101925 | 658 | 15 |
| 65 | BC ZSM Lep 101926 | 658 | 15 |
| 66 | BC ZSM Lep 101927 | 658 | 15 |
| 67 | BC ZSM Lep 101928 | 658 | 17 |
| 68 | BC ZSM Lep 101929 | 658 | 17 |
| 69 | BC ZSM Lep 101930 | 658 | 17 |
| 70 | BC ZSM Lep 101931 | 658 | 18 |
| 71 | BC ZSM Lep 101932 | 658 | 19 |
| 72 | BC ZSM Lep 101933 | 658 | 19 |
| 73 | BC ZSM Lep 101934 | 658 | 19 |
| 74 | BC ZSM Lep 101935 | 658 | 19 |
| 75 | BC ZSM Lep 101936 | 658 | 19 |
| 76 | BC ZSM Lep 101937 | 658 | 19 |
| 77 | BC ZSM Lep 101938 | 658 | 19 |
| 78 | BC ZSM Lep 101939 | 658 | 20 |
| 79 | BC ZSM Lep 101940 | 658 | 20 |
| 80 | BC ZSM Lep 101941 | 658 | 20 |
| 81 | BC ZSM Lep 101942 | 658 | 20 |
| 82 | BC ZSM Lep 101943 | 658 | 21 |
| 83 | BC ZSM Lep 101944 | 658* | 21 |
| 84 | BC ZSM Lep 101945 | 658 | 22 |
| 85 | BC ZSM Lep 101946 | 658 | 23 |
| 86 | BC ZSM Lep 101947 | 658 | 23 |
| 87 | BC ZSM Lep 101948 | 0 | 23 |
| 88 | BC ZSM Lep 101949 | 658 | 23 |
| 89 | BC ZSM Lep 101950 | 658 | 23 |
| 90 | BC ZSM Lep 101951 | 658 | 23 |
| 91 | BC ZSM Lep 101952 | 658 | 23 |
| 92 | BC ZSM Lep 101953 | 658 | 23 |
| 93 | BC ZSM Lep 101954 | 658 | 24 |
| 94 | BC ZSM Lep 101955 | 0 | 25 |
| 95 | BC ZSM Lep 101956 | 658 | 25 |
| 96 | BC ZSM Lep 101957 | 658 | 27 |
| 97 | BC ZSM Lep 101958 | 658 | 27 |
| 98 | BC ZSM Lep 101959 | 658 | 28 |
| 99 | BC ZSM Lep 101960 | 658 | 29 |
| 100 | BC ZSM Lep 101961 | 658 | 29 |
| 101 | BC ZSM Lep 101962 | 658 | 32 |
| 102 | BC ZSM Lep 101963 | 658* | 32 |
| 103 | BC ZSM Lep 101964 | 658 | 32 |
| 104 | BC ZSM Lep 101965 | 658 | 32 |
| 105 | BC ZSM Lep 101966 | 658 | 32 |
| 106 | BC ZSM Lep 101967 | 0 | 32 |
| 107 | BC ZSM Lep 101968 | 658* | 32 |
| 108 | BC ZSM Lep 101969 | 658* | 32 |
| 109 | BC ZSM Lep 101970 | 658 | 32 |
| 110 | BC ZSM Lep 101971 | 658 | 32 |
| 111 | BC ZSM Lep 101972 | 658 | 33 |
| 112 | BC ZSM Lep 101973 | 658 | 33 |
| 113 | BC ZSM Lep 101974 | 658 | 34 |
| 114 | BC ZSM Lep 101975 | 658 | 35 |
| 115 | BC ZSM Lep 101976 | 658 | 35 |
| 116 | BC ZSM Lep 101977 | 658 | 36 |
| 117 | BC ZSM Lep 101978 | 658 | 36 |
| 118 | BC ZSM Lep 101979 | 0 | 36 |
| 119 | BC ZSM Lep 101980 | 658 | 37 |
| 120 | BC ZSM Lep 101981 | 658 | 37 |
| 121 | BC ZSM Lep 101982 | 658 | 38 |
| 122 | BC ZSM Lep 101983 | 658 | 38 |
| 123 | BC ZSM Lep 101984 | 658 | 40 |
| 124 | BC ZSM Lep 101985 | 0 | 40 |
| 125 | BC ZSM Lep 101986 | 0 | 40 |
| 126 | BC ZSM Lep 101987 | 658 | 41 |
| 127 | BC ZSM Lep 101988 | 658 | 42 |
| 128 | BC ZSM Lep 101989 | 658 | 44 |
| 129 | BC ZSM Lep 101990 | 0 | 44 |
| 130 | BC ZSM Lep 101991 | 658 | 47 |

Identity of larva see Table 2, identity of trees see Table 3 and Supporting information S1+S3
Table; \* sequenced by AIM company with special primers for alcaloid-inhibited samples.

**Table 2.** Identification results from sequence blasting on BOLD for 119 successfully sequenced Lepidoptera larvae (see Table 1) and their distances from the nearest genetic neighbour.

| Larva Nr. | Family | Identification of larvae from BOLD-blast | Nearest neighbour (NN) on BOLD | distance from NN (%) | category of match |
| --- | --- | --- | --- | --- | --- |
| 1 | Bombycidae | *Quentalia* | *Quentalia chromana*DHJ01 | 5.8 | genus |
| 2 | Bombycidae | *Quentalia* | *Quentalia chromana*DHJ01 | 5.8 | genus |
| 3 | Erebidae | *Lascoria* | *Lascoria* species indet. | 0.45 | species |
| 4 | Gelechiidae | Gelechiidae | Gelechiidae genus indet. | 8.0 | family |
| 5 | Gelechiidae | *Dichomeris* | *Dichomeris* species indet. | 4.6 | genus |
| 6 | Geometridae | Larentiinae | *Xanthorhoe labradorensis* | 8.1 | family |
| 7 | Plutellidae | Plutellidae | *Rhigognostis senilella* | 8.9 | family |
| 8 | Erebidae | *Ypsora selenodes* | *Ypsora selenodes* | 1.2 | species |
| 9 | Geometridae | *Thysanopyga apicitruncaria* | *Thysanopyga apicitruncaria* | 0.15 | species |
| 10 | Tineidae | *Hybroma* | *Hybroma* species indet. | 6.9 | genus |
| 11 | Crambidae | Spilomelinae_genus sp. 30YB | Spilomelinae_genus sp. 30YB | 1.4 | species |
| 12 | Crambidae | Spilomelinae_genus sp. 30YB | Spilomelinae_genus sp. 30YB | 1.1 | species |
| 13 | Geometridae | *Hemipterodes divaricata* | *Hemipterodes* BioLep111 | 3.5 | genus |
| 14 | Erebidae | Erebidae | *Bertula tespisalis* | 6.6 | family |
| 15 | Erebidae | Erebidae | *Bertula tespisalis* | 6.6 | family |
| 16 | Erebidae Cte. | *Delphyre orientalis* | *Delphyre orientalis* | 1.2 | species |
| 17 | Riodinidae | Riodinidae | *Semomesia croesus* | 7.5 | family |
| 18 | Erebidae Lit. | *Prepiella* species 1 | *Prepiella* species indet. | 0.15 | species |
| 19 | Gelechiidae | Gelechiidae | Gelechiidae genus indet. | 2.4 | genus |
| 20 | Geometridae | *Patalene hamulata*AH01Pe | *Patalene hamulata*AH01Pe | 0.0 | species |
| 21 | Depressariidae | Depressariidae | *Antaeotricha Janzen*86 | 7.5 | family |
| 22 | Uraniidae | Uraniidae species indet. | Uraniidae species indet. | 0.0 | species |
| 23 | Uraniidae | Uraniidae species indet. | Uraniidae species indet. | 0.0 | species |
| 24 | Noctuidae | noctBioLep01 BioLep2008 | noctBioLep01 BioLep2008 | 1.4 | species |
| 26 | Uraniidae | *Urania leilus* | *Urania leilus* | 0.0 | species |
| 27 | Depressariidae | Depressariidae | *Stenoma* species indet. | 7.9 | family |
| 28 | Depressariidae | Depressariidae | *Stenoma* species indet. | 7.9 | family |
| 29 | Uraniidae | Uraniidae | *Cyphyra swinhoi* (Uran.) | 7.2 | family |
| 30 | Erebidae Lit. | *Nodozana* nr. *coresa* | *Nodozana* nr. *coresa* | 0.15 | species |
| 31 | Gelechiidae | Gelechiidae | GelJanzen01 Janzen180 | 7.8 | family |
| 32 | Apatelodidae | *Olceclostera* | *Olceclostera* species indet. | 6.9 | genus |
| 33 | Depressariidae | Depressariidae | *Stenoma* species indet. | 6.9 | family |
| 34 | Depressariidae | Depressariidae | *Stenoma* species indet. | 6.7 | family |
| 35 | Gelechiidae | Dichomeris? | Gelechiidae genus indet. | 2.2 | species |
| 36 | Depressariidae | Depressariidae | Depressariidae genus indet. | 2.6 | genus |
| 37 | Noctuidae | Noctuidae_incertae_sedis sp. 14YB | Noctuidae_incertae_sedis sp. 14YB | 1.9 | species |
| 38 | Geometridae | *Ergavia* | *Ergavia* species indet. | 3.8 | genus |
| 39 | Notodontidae | Notodontidae | *Hemiceras plana* | 9.4 | family |
| 40 | Geometridae | *Physocleora* AH02Pe | *Physocleora* AH02Pe | 0.15 | species |
| 42 | Bombycidae | *Quentalia* | *Quentalia chromana*DHJ01 | 5.8 | genus |
| 43 | Hesperiidae | *Panoquina fusina* | *Panoquina fusina* | 0.0 | species |
| 45 | Gelechiidae | Gelechiidae | Gelechiidae genus indet. | 8.5 | family |
| 46 | Geometridae | Larentiinae species | Larentiinae genus indet. | 0.15 | species |
| 47 | Erebidae | *Mecodina* | *Mecodina* species indet. | 8.1 | genus |
| 48 | Apatelodidae | *Olceclostera* | *Olceclostera* species indet. | 6.4 | genus |
| 49 | Erebidae | *Deinopa* | *Deinopa angitia* | 5.1 | genus |
| 50 | unidentified | Lepidoptera | *Semomesia croesus* (Riodin.) | 9.6 | order |
| 51 | unidentified | Lepidoptera | *Semomesia croesus* (Riodin.) | 9.8 | order |
| 52 | Erebidae | *Eudocima procus* | *Eudocima procus* | 0.0 | species |
| 53 | Erebidae | *Gorgone umbrigens*DHJ02 | *Gorgone umbrigens*DHJ02 | 0.6 | species |
| 54 | unidentified | Lepidoptera | *Semomesia croesus* (Riodin.) | 9.6 | order |
| 55 | Crambidae | *Evergestis* | *Evergestis simulatilis* | 6.1 | genus |
| 56 | Erebidae | Erebidae | *Catocala relecta* | 6.6 | family |
| 58 | Geometridae | *Perissopteryx divisaria* | *Perissopteryx divisaria* | 1.4 | species |
| 59 | Geometridae | *Thysanopyga apicitruncaria* | *Thysanopyga apicitruncaria* | 0.15 | species |
| 60 | Erebidae | *Lascoria* Poole03 | *Lascoria* Poole03 | 0.45 | species |
| 61 | Erebidae | *Letis magna* | *Letis magna* | 0.0 | species |
| 62 | Saturniidae | *Automeris denticulata* | *Automeris denticulata* | 0.0 | species |
| 63 | Bombycidae | *Anticla* | *Anticla antica*DHJ04 | 2.9 | genus |
| 64 | Erebidae | *Eudocima procus* | *Eudocima procus* | 0.0 | species |
| 65 | Erebidae | *Eudocima procus* | *Eudocima procus* | 0.0 | species |
| 66 | Erebidae Phae. | *Ernassa* species | *Ernassa* species indet. | 2.1 | species |
| 67 | Phiditiidae | *Phiditia* | *Phiditia lucernaria* | 3.8 | genus |
| 68 | Geometridae | *Pero incisa* | *Pero incisa* | 0.15 | species |
| 69 | Erebidae | *Lascoria* Poole03 | *Lascoria* Poole03 | 0.45 | species |
| 70 | Erebidae | *Metalectra* | *Metalectra* BioLep167 | 5.2 | genus |
| 71 | Erebidae | *Latebraria amphipyroides* | *Latebraria amphipyroides* | 0.6 | species |
| 72 | Noctuidae | *Drobeta* | *Drobeta* Poole17 | 4.1 | genus |
| 73 | Erebidae | *Latebraria amphipyroides* | *Latebraria amphipyroides* | 0.6 | species |
| 74 | Geometridae | *Idaea orilochia* | *Idaea orilochia* | 0.0 | species |
| 75 | Geometridae | *Hemipterodes divaricata* | *Hemipterodes* BioLep111 | 3.5 | genus |
| 76 | Apatelodidae | Apatelodidae | Apatelodidae genus indet. | 6.6 | family |
| 77 | Erebidae | *Mastixis* | *Mastixis* Poole02 | 3.5 | genus |
| 78 | Erebidae | *Feigeria scops* | *Feigeria scops* | 0.0 | species |
| 79 | Geometridae | Sterrhinae | *Cyclophora* species indet. | 6.1 | family |
| 80 | Geometridae | *Semaepopus* | *Semaepopus* Janzen216 | 5.5 | genus |
| 81 | Nymphalidae | *Memphis acidalia* | *Memphis acidalia* | 0.0 | species |
| 82 | Euteliidae | *Paectes* | *Paectes circularis* | 3.7 | genus |
| 83 | Erebidae Cte. | *Calonotos chalcipleura* (Ereb.) | *Calonotos chalcipleura* | 0.0 | species |
| 84 | Erebidae Lit. | *Clemensia* | Erebidae genus indet. (*Clemensia*) | 1.7 | species |
| 85 | Erebidae Lit. | *Prepiella* species 2 | *Prepiella* species indet. | 0.3 | species |
| 86 | Geometridae | *Patalene hamulata*AH01Pe | *Patalene hamulata*AH01Pe | 0.0 | species |
| 88 | Hesperiidae | *Polythrix* | *Polythrix kanshul* | 5.3 | genus |
| 89 | Erebidae Phae. | *Stidzaeras strigifera* | *Stidzaeras strigifera* | 2.2 | species |
| 90 | Erebidae Lit. | *Apistosia judas* | *Apistosia judas* | 0.0 | species |
| 91 | Erebidae Phae. | *Stidzaeras strigifera* | *Stidzaeras strigifera* | 1.9 | species |
| 92 | Saturniidae | *Pseudautomeris arminirene* | *Pseudautomeris arminirene* | 0.8 | species |
| 93 | Uraniidae | Uraniidae species indet. | Uraniidae species indet. | 0.0 | species |
| 95 | Erebidae | *Lascoria manes* | *Lascoria manes* | 0.8 | species |
| 96 | Erebidae | *Clapra* | *Clapra* species indet | 2.1 | species |
| 97 | Erebidae | *Lascoria* Poole03 | *Lascoria* Poole03 | 0.0 | species |
| 98 | Saturniidae | *Homoeopteryx* | *Homoeopteryx major* | 2.7 | genus |
| 99 | Erebidae Cte. | *Haemanota nigricollum* | *Haemanota nigricollum* | 0.0 | species |
| 100 | Erebidae Cte. | *Haemanota nigricollum* | *Haemanota nigricollum* | 0.0 | species |
| 101 | Notodontidae | *Kaseria* | *Kaseria pallida* | 4.6 | genus |
| 102 | Geometridae | *Mychonia* (Geom.) | *Mychonia* | 0.0 | species |
| 103 | Noctuidae | *Lycaugesia* | Noctuidae genus indet. (*Lycaugesia*) | 3.2 | genus |
| 104 | Geometridae | *Ischnopteris chlorophaearia* | *Ischnopteris chlorophaearia* | 0.0 | species |
| 105 | Apatelodidae | *Olceclostera* | *Olceclostera* species indet. | 2.6 | genus |
| 107 | Erebidae | *Lascoria* Poole03 (Ereb.) | *Lascoria* Poole03 | 0.0 | species |
| 108 | Erebidae Lit. | *Apistosia judas* (Ereb.) | *Apistosia judas* | 0.0 | species |
| 109 | Nymphalidae | *Euptychia* n. sp. 5 CP-2006 | *Euptychia* n. sp. 5 CP-2006 | 1.6 | species |
| 110 | Apatelodidae | Apatelodidae species | Apatelodidae genus indet. | 2.0 | species |
| 111 | Erebidae | *Antiblemma sterope* | *Antiblemma sterope* | 0.5 | species |
| 112 | Erebidae | *Eudocima* | *Eudocima* species indet. | 6.1 | genus |
| 113 | Bombycidae | *Anticla* | *Anticla antica*DHJ03 | 6.6 | genus |
| 114 | Erebidae | *Sosxetra grata* | *Sosxetra grata* | 0.0 | species |
| 115 | Hesperiidae | *Myscelus epimachia* | *Myscelus epimachia* | 0.0 | species |
| 116 | Geometridae | *Glena* AH03Pe | *Glena* AH03Pe | 0.15 | species |
| 117 | Geometridae | *Semiothisa gambaria* | *Semiothisa gambaria* | 0.0 | species |
| 119 | Erebidae Phae. | *Melese* | *Melese drucei* | 3.2 | genus |
| 120 | Geometridae | *Physocleora* AH02Pe | *Physocleora* AH02Pe | 0.3 | species |
| 121 | Geometridae | *Stegotheca* | *Stegotheca* species indet. | 3.5 | genus |
| 122 | Erebidae Phae. | *Pelochyta arontes* | *Pelochyta arontes* | 0.0 | species |
| 123 | Geometridae | *Stegotheca* | *Stegotheca* species indet. | 3.2 | genus |
| 126 | Erebidae | *Sosxetra grata* | *Sosxetra grata* | 0.0 | species |
| 127 | Noctuidae | Noctuidae | Noctuidae/Acontiinae genus indet. | 4.0 | genus |
| 128 | Erebidae | Erebidae | *Phytometra ernestinana* | 8.5 | family |
| 130 | Geometridae | *Stegotheca* | *Stegotheca* species indet. | 3.5 | genus |

Lit. = Lithosiini (Arctiinae); Cte. = Ctenuchina (Arctiinae); Phae. = Phaegopterina (Arctiinae)

**Table 3.** Identification of target trees, results from blasting on NCBI (BLAST matches usually >99%).

| Target tree nr. | Preidentification (from vernacular names; cf. Supporting info. S1) | Molecular consensus identification (rbcL, trnL-F and psbA genes; cf. Supp. info. S3) | Consensus identification |
| --- | --- | --- | --- |
| 1 | *Mangifera indica* | *Mangifera indica* | *Mangifera indica* (Anacardiaceae) |
| 2 | *Mangifera indica* | *Mangifera indica* | *Mangifera indica* (Anacardiaceae) |
| 3 | Meliaceae or Annonaceae | *Guarea* or *Cabrlea* (Meliaceae) | *Guarea* or *Cabrlea* (Meliaceae) |
| 4 # | Anacardiaceae | *Mangifera* or *Spondias* (Anacardiaceae) | *Mangifera* or *Spondias* (Anacardiaceae) |
| 5 | *Guarea* (Meliaceae) | *Guarea* or *Cabrlea* (Meliaceae) | *Guarea* (Meliaceae) |
| 6 # | *Ficus* (Moraceae) | *Ficus* (Moraceae) | *Ficus* (Moraceae) |
| 7 | ,Ucu muchaca' <sup>1</sup> | Malvaceae or Meliaceae <sup>2</sup> | Malvaceae or Meliaceae <sup>2</sup> |
| 8 # | *Apeiba* (Malvaceae) | Malvaceae | *Apeiba* (Malvaceae) |
| 9 | *Leonia glycyarpa* (Violaceae) | *Leonia glycyarpa* (Violaceae) | *Leonia glycyarpa* (Violaceae) |
| 10 | Annonaceae | *Oxandra polyantha* (Annonaceae) | *Oxandra polyantha* (Annonaceae) |
| 11-1 | *Celtis schippii* (Cannabaceae) | *Celtis schippii* (Cannabaceae) | *Celtis schippii* (Cannabaceae) |
| 11-2 | *Neea* (*Guapira*) (Nyctaginaceae) | *Neea* (Nyctaginaceae) | *Neea* (Nyctaginaceae) |
| 12 | Annonaceae | *Oxandra polyantha* (Annonaceae) and/or *Conceveiba guianensis* (Euphorbiaceae) <sup>2</sup> | *Oxandra polyantha* (Annonaceae) and/or *Conceveiba guianensis* (Euphorbiaceae) <sup>2</sup> |
| 13 | Annonaceae | *Oxandra polyantha* (Annonaceae) | *Oxandra polyantha* (Annonaceae) |
| 14 | *Poulsenia armata* (Moraceae) | *Naucleopsis* (Moraceae) | *Poulsenia* or *Naucleopsis* (Moraceae) |
| 15 | ,Ucu muchaca' <sup>1</sup> | *Hirtella* (Chrysobalanaceae) | *Hirtella* (Chrysobalanaceae) |
| 16 | *Castilla* | *Castilla elastica* (Moraceae) | *Castilla elastica* (Moraceae) |
| 17 | Moraceae | *Clarisia biflora* (Moraceae) | *Clarisia biflora* (Moraceae) |
| 18 | *Ficus* (Moraceae) | *Ficus* (Moraceae) | *Ficus* (Moraceae) |
| 19 | Annonaceae | *Oxandra polyantha* (Annonaceae) | *Oxandra polyantha* (Annonaceae) |
| 20 | no name provided | *Neea* (Nyctaginaceae) | *Neea* (Nyctaginaceae) |
| 21 | *Apeiba* (Malvaceae) | *Annona* (Annonaceae) | *Annona* (Annonaceae) or *Apeiba* sp. (Malvaceae) |
| 22 | ,Kaimitio' | *Byrsonima coccolobifolia* (Malpighiaceae) | *Byrsonima coccolobifolia* (Malpighiaceae) |
| 23 | no name provided | no tissue provided | unidentified |
| 24 | *Perebea* (Moraceae) | *Pouteria* or *Chrysophyllum* (Sapotaceae) | Moraceae or Sapotaceae <sup>3</sup> |
| 25 | *Otoba parvifolia* (Myristicaceae) | Myristicaceae | *Otoba parvifolia* (Myristicaceae) |
| 26 # | *Apeiba* (Malvaceae) | Malvaceae | *Apeiba* (Malvaceae) |
| 27 | Annonaceae | *Oxandra polyantha* (Annonaceae) | *Oxandra polyantha* (Annonaceae) |
| 28 | *Ficus* (Moraceae) | *Ficus* (Moraceae) and/or *Simira* (Rubiaceae) <sup>2</sup> | *Ficus* (Moraceae) and/or *Simira* (Rubiaceae) <sup>2</sup> |
| 29 | *Apeiba* (Malvaceae) | Malvaceae | *Apeiba* (Malvaceae) |
| 30 # | *Garcinia* (Clusiaceae) | *Garcinia macrophylla* or *G. mangostana* (Clusiaceae) | *Garcinia macrophylla* or *G. mangostana* (Clusiaceae) |
| 31-1 # | *Garcinia* (Clusiaceae) | *Garcinia* (Clusiaceae) | *Garcinia* (Clusiaceae) |
| 31-2 # | ,Tawari' | Sapotaceae or Fabaceae <sup>2</sup> | Sapotaceae or Fabaceae <sup>2</sup> |
| 32 | Sapindaceae | *Paullinia* (Sapindaceae) | *Paullinia* (Sapindaceae) |
| 33 | *Guarea* (Meliaceae) <sup>1</sup> | *Trichilia* (Meliaceae) | *Trichilia* (Meliaceae) |
| 34 | Moraceae | *Ficus* (Moraceae) | *Ficus* (Moraceae) |
| 35 | *Guarea* (Meliaceae) <sup>1</sup> | *Guarea guidonia* (Meliaceae) | *Guarea guidonia* (Meliaceae) |
| 36 | *Guarea* (Meliaceae) <sup>1</sup> | *Guarea guidonia* (Meliaceae) | *Guarea guidonia* (Meliaceae) |
| 37 | *Guarea* (Meliaceae) <sup>1</sup> | *Guarea guidonia* (Meliaceae) | *Guarea guidonia* (Meliaceae) |
| 38 | *Guarea* (Meliaceae) <sup>1</sup> | *Erythrina speciosa* (Fabaceae) <sup>2</sup> | *Guarea* (Meliaceae) or *Erythrina* (Fabaceae) <sup>1 2</sup> |
| 39 # | *Tapirira guianensis* (Anacardiaceae) | *Tapirira guianensis* (Anacardiaceae) or *Guarea guidonia* (Meliaceae) <sup>1 2</sup> | *Tapirira guianensis* (Anacardiaceae) or *Guarea guidonia* (Meliaceae) <sup>1 2</sup> |
| 40 | *Guarea* (Meliaceae) <sup>1</sup> | *Guarea guidonia* (Meliaceae) | *Guarea guidonia* (Meliaceae) |
| 41 | *Guarea* (Meliaceae) <sup>1</sup> | *Guarea guidonia* (Meliaceae) | *Guarea guidonia* (Meliaceae) |
| 42 | *Ficus* (Moraceae) | *Ficus* (Moraceae) | *Ficus* (Moraceae) |
| 43 # | *Guarea* (Meliaceae) <sup>1</sup> | *Guarea guidonia* (Meliaceae) | *Guarea guidonia* (Meliaceae) |
| 44 | *Guarea* (Meliaceae) <sup>1</sup> | *Guarea guidonia* (Meliaceae) | *Guarea guidonia* (Meliaceae) |
| 45 # | *Guarea* (Meliaceae) <sup>1</sup> | *Guarea guidonia* (Meliaceae) | *Guarea guidonia* (Meliaceae) |
| 46 # | *Guarea* (Meliaceae) <sup>1</sup> | *Guarea guidonia* (Meliaceae) | *Guarea guidonia* (Meliaceae) |
| 47 | *Guarea* (Meliaceae) <sup>1</sup> | *Guarea guidonia* (Meliaceae) | *Guarea guidonia* (Meliaceae) |

**Fig 1.**
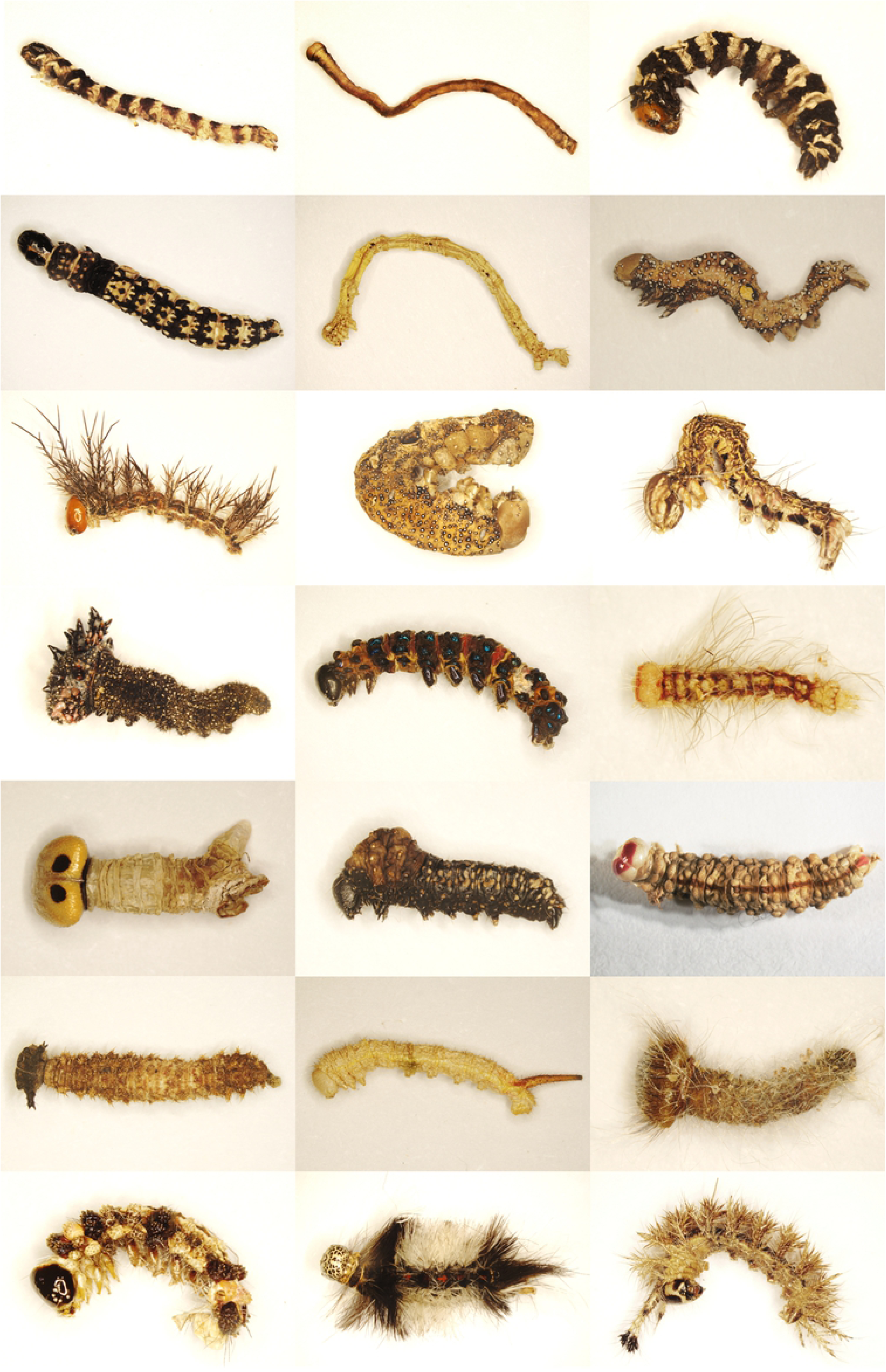
Lepidopteran larvae after selection from the alcohol-preserved fogging samples from Panguana, Peru, after drying and before tissue-sampling for the DNA analysis. (Upper) row 1: larvae nr. 5, 13, 26; row 2: nr. 28, 46, 52; row 3: nr. 62, 65, 71; row 4: nr. 81, 83, 85; row 5: nr. 88, 93, 100; row 6: nr. 109, 113, 115; (bottom) row 7: nr. 119, 122, 124 (numbers and identification of larvae see Table 2).

Tissue samples were submitted to the standard procedures of the Canadian Centre for DNA Barcoding (CCDB) for sequencing the mitochondrial 5’ cytochrome oxidase gene, subunit 1 (COI), the standard marker for the identification of most animals. LepF1 and LepR1 were the primers used for PCR and sequencing [28]. Sequences were blasted against the complete sequence database of the Barcode of Life Data systems (BOLD, [29]) in order to infere the closest matches using the BOLD Identification Engine (http://www.boldsystems.org/index.php/IDS_OpenIdEngine). Also morphology of larvae and related (genetically near) adult moths were considered to test the reliability of the results. Nomenclature of scientific taxon names follows the catalogue used on BOLD database, which in many families is in accordance with the currently available catalogues (e.g. [30] for Geometridae). Vouchers of larvae are stored at the Zoologische Staatssammlung München, Germany. Sequences, images and related metadata are available open access on BOLD under the dataset DS-PANLARVA (dx.doi.org/10.5883/DS-PANLARVA).

### Tissue sampling and morphology-based identification of target trees

The 47 target trees have been pre-identified in the field based on morphology (shape of tree growth and shape of leaves, rarely blossoms or fruits) by the native caretaker of the Panguana Station, “Moro” Carlos Vásquez Módena, to Peruvian vernacular names (see Table 3 and Supporting information S1 Table), which usually cannot be unequivocally referred to scientific plant names, however. For a tentative assignment of vernacular names to botanical taxa see Supporting information S1 Table. For nomenclature of plant names we follow the “Plant List” (available online at www.theplantlist.org/1/). For most target trees a small branch was collected, pressed and kept in a herbarium for identification. Identification of a selection of sampled leaves was performed by Hamilton Paredes, Museo de Historia Natural, Lima. A small leaf piece was cut as tissue sample for DNA-Barcoding. In addition to that, sapwood/cambium tissue samples were taken of each target tree by using a leather punch to extract a core from the stem. Then, a thin slice of sapwood/cambium was cut and immediately dried over silica gel.

### Identification of target trees through DNA barcoding (trnL-F, rbcL & psbA)

Because of the above mentioned uncertainties of target tree pre-identification, we have submitted plant tissue samples to DNA barcoding. For that purpose leaves were available for 37 out of the 47 target trees, pieces of cambium+sapwood for 46 trees. Plant tissues (leaves) were submitted to Sanger sequencing (AIM; Advanced Identification Methods GmbH – www.aimethods-lab.com) with two markers, rbcL and psbA using standardized protocols following [31], [32]. An additional attempt was performed in CCDB (Guelph, Canada; primers: trnL-F, rbcL; standard Sanger sequencing procedure) using both leaves and sapwood samples, the latter supplementing those cases where no leaves were available for study. All resulting sequences were blasted against GenBank (NBCI) and BOLD data using standard blast functions. Sequences and related metadata are available open access on BOLD under the dataset DS-PANPLANT (dx.doi.org/10.5883/DS-PANPLANT).

### Gut content analysis (rbcL, psbA, ITS2, matK)

For a subset of ten larvae, gut content analysis was tested for molecular identification of the larva’s ‘true’ diet. For that purpose, we performed a second vertical cut and submitted one segment of the larva to High-Throughput-Sequencing (HTS) with the two markers rbcL and psbA. Cut slices of the caterpillars were dried, homogenized and DNA extracted using the DNEasy Plant kit (Qiagen, Hilden, Germany). HTS-adapted versions of the target gene primers were used for PCR amplification. Successfully amplified products were used for a subsequent PCR reaction which adds Illumina Nextera XT indices to each PCR product, enabling a unique tagging of each sample. Indexed samples were measures using a Qubit v4.0 (Life Technologies) and combined into equilmolar pools, which were then size selected using preparative gel electrophoresis. Gel-extracted pools were then measured again and combined in the final library, which was sequenced using v.2 chemistry on an Illumina MiSeq (2*250bp).

## Results

### Identification of larvae

A total of 130 caterpillar specimens were collected from 37 of the 47 target plants. In the samples of 10 target trees lepidopteran larvae were not found. COI sequencing (DNA barcoding) was successful for 119 larvae (91.5%). The larvae belong to 92 different COI clusters (BINs), which are a good proxy for different species [33].

When blasting the DNA barcodes of the larvae on BOLD database, 65 larvae (55%) belonging to 48 species showed ‘close genetic similarity’ – here defined as lower than 2.5% – with adult reference vouchers. Such genetic similarity is interpreted here as ‘species (or sister species) level matches’ (Table 2). 27 species have Linnean names on BOLD database, 20 are listed under ‘interim names’ (name codes) which either refer to described but not-yet-identified taxa or to undescribed species.

For 32 larvae (27%) belonging to 27 species the blasting on BOLD database revealed genus level matches, in five cases with disputable reliability. For 19 larvae (16%) assignment to subfamily or family level was possible, the reliability of 12 of these assignments needs to be tested by further extension of the reference database, since long branch attraction effects may have influenced the results in a few single cases. In just three cases belonging to one single species no family suggestion could be given based on the COI barcode.

### Identification of target plants

Pre-identification of target trees, as performed by the local administrator of the Panguana station (see Supporting information S1 Table), was supplemented by molecular identification (sequencing of leaves and sapwood with the markers trnL-F, rbcL and psbA) of all but one of the target trees. For all but one of the target trees (98%) molecular identification through blasting on BOLD and GenBank brought a reliable identification to at least family level (see Table 3 and Supporting information S3 Table) but in four cases two different families resulted which apparently is due to sampling of leaves and sapwood from different plants. In 37 target trees (79%) identification was possible to genus or species level (see Table 3 and Supporting information S3 Table).

### Gut content analysis

Gut content analysis was performed for ten larvae based on Next-Generation-Sequencing with two markers rbcL and psbA. The two highest numbers of HTS-reads for rbcL and psbA genes and their genetically most similar species as resulting from BLAST-search in GenBank is shown for each larva in Table 4.

**Table 4.** Gut contents of ten fogged larvae with identity of target tree and HTS results from molecular identification of gut content, only the BLAST identification of the fragements with the two most numerous reads shown.

| Nr. and identity of larva | Nr. and identity of target tree | HTS gut content (best hit): *r*(*bcl*), *p*(*sbA*) | nr. of reads | HTS gut content (second best hit) | nr. of reads |
| --- | --- | --- | --- | --- | --- |
| 40 *Physocleora* AH02Pe (Geom.) | 2 *Mangifera indica* (Anac.) | *Mangifera indica* (Anac.) *r*+*p* | 12076 | *Toxicodendron pubescens* (Anac.) *r* | 1115 |
| 42 *Quentalia* (Bomb.) | 3 *Guarea* or *Cabralea* (Meli.) | *Trophis racemosa* (Mora.) *r* | 27876 | (contaminations) | 1-526 |
| 65 *Eudocima procutus* (Ereb.) | 15 *Hirtella* (Chrys.) | *Tinospora smilacina* (Meni.) <sup>1</sup> *r* | 92678 | *Odontocarya tamoides* (Meni.) <sup>1</sup> *r* | 78516 |
| 76 Apatelodidae | 19 *Oxandra polyantha* (Anno.) | *Cucumis sativus* (Cucu.) *r* | 39129 | *Lasthenia californica* (Aste.) *r* | 37725 |
| 82 *Paectes* nr *circularis* (Noct.) | 21 *Annona* (Anno.) or *Apeiba* (Malv.) | *Lejeunea bidentula* (Bryo.) *p* | 6196 | *Lejeunea tuberculosa* (Bryo.) *p* | 5967 |
| 83 - *Colanotos chalcipleura* (Ereb.) | 21 *Annona* (Anno.) or *Apeiba* (Malv.) | *Echites yucatanensis* (Apoc.) <sup>2</sup> *p* | 73311 | *Anodendron* cf. *affine* (Apoc.) <sup>3</sup> *p* | 68464 |
| 98 *Homoeopteryx* nr *major* (Satu.) | 28 *Ficus* (Mora.) and/or *Simira* (Rubi.)! | *Faramea occidentalis* (Rubi.) *r* | 109687 | *Faramea pedunculata* (Rubi.) *r* | 92548 |
| 102 - *Mychonia* (Geom.) | 32 *Paullinia* (Sapi.) | *Ceratolejeunea diversicornua* (Bryo.) <sup>5</sup> *p* | 925060 | *Schizocolea linderi* (Rubi.) <sup>4</sup> *p* | 64812 |
| 107 - *Lascaria* Poole03 (Ereb.) | 32 *Paullinia* (Sapi.) | *Lejeunea bidentula* (Bryo.) <sup>5</sup> *p* | 1550587 | *Schizocolea linderi* (Rubi.) <sup>4</sup> *p* | 395941 |
| 108 - *Apistosia judas* (Ereb.) | 32 *Paullinia* (Sapi.) | *Schizocolea linderi* (Rubi.) <sup>4</sup> *p* | 934090 | *Nyholmiella obtusifolia* (Bryo.) *p* | 482948 |

*p* = *psbA*; *r* = *rbcL*. <sup>1</sup> liana, Australian; - <sup>2</sup> liana, South American; - <sup>3</sup> Asian; - <sup>4</sup> African; - <sup>5</sup> with several other sub-optimal blast hits in Bryophyta. Abbreviations of lepidopteran families: Geom. = Geometridae; Bomb. = Bombycidae; Ereb. = Erebidae; Noct. = Noctuidae; Satu. = Saturniidae. Abbreviations of plant families: Anac. = Anacardiaceae; Mora. = Moraceae; Meli. = Meliaceae; Chrys. = Chrysobalanaceae; Anno. = Annonaceae; Malv. = Malvaceae; Rubi. = Rubiaceae; Sapi. = Sapindaceae; Meni. = Menispermaceae; Cucu. = Cucurbitaceae; Bryo. = Bryophyta; Apoc. = Apocynaceae; Aste. = Asteraceae.

## Discussion

When investigating host-plant relationships it is usually assumed that larvae feed on the plants from where they have been collected. This assumption is based on the behaviour of larvae usually resting on their feeding plant during their development. One needs to consider, however, that certain larvae abandon their host-plants searching for a hidden resting place during daytime and mature larvae often leave their food-plant in the last days before pupation looking for a suitable pupation site, sometimes far from their feeding plants. Moreover, in particular in rainwood forests “alternative feeders” may use epiphytes, lianas, lichens, algae, fungi or mosses [13], [14], and in our fogging approach pitfalls are possible through collateral fogging of larvae from neighboring trees. Gut content analysis can shed light on true feeding biology.

### Gut content matching identity of target tree

Only in one out of ten analysed larvae (see Table 4) the gut content revealed to match exactly the fogged target tree species: *Physocleora* AH02Pe (Geometridae; larva nr. 40) fogged from *Mangifera indica*. In a second case, *Homoeopteryx* near *major* (Saturniidae; larva nr. 98), the gut content revealed to be from the same plant family (Rubiaceae), genus *Simira* resulting from sequencing of plant sapwood and genus *Faramea* resulting from gut content analysis. The high percentage of eight out of ten larvae with a mismatch between target tree and gut content suggests that an a priori assignation of fogged larvae to the target trees usually is erroneous and that alternative feeding (epiphytes, algae, mosses etc.) or feeding on lianas and neighboring trees plays a major role. The rate of alternative feeding should be tested basing on a larger sample, ruling out a potentially biased ratio through external contamination of larvae by plant DNA (see below).

### Gut content matching previously known host-plant but not the target tree

A larva of the genus *Quentalia* (Bombycidae, larva nr. 42, see Table 4) was fogged from a tree of the family Meliaceae, but the gut content pointed to feeding on *Trophis racemosa* (Moraceae). Since *Quentalia* larvae were previously recorded as feeding on Moraceae [4], *Trophis racemosa* is likely the true food-plant of the *Quentinalia* larva which may have been growing close to the target tree.

In a second case, *Eudocima procus* (Erebidae; larva nr. 65, see Table 4) was fogged from a tree of the genus *Hirtella* (Chrysobalanaceae) but the gut content pointed to feeding on *Tinospora smilacina* (Menispermaceae). Since species of the genus *Eudocima* are known to feed on Menispermaceae [34], [35], [4], *Tinospora smilacina* is likely the true food-plant of the *Eudocima* larva. *Tinospora* is a liana and likely was associated with the target tree. A similar case is also referring to liana-feeding: larva nr. 83 (see Table 4) was fogged from a tree of Annonaceae or Malvaceae, but in its gut content we found the DNA of the neotropical liana *Echites yucatanensis* (Apocynaceae).

Hence in three out of ten cases (30%) feeding on lianas or on a neighboring tree was recorded. Although the rate of feeding on such associated or neighbouring plants should be tested basing on a larger sample, the results of this pilot study clearly show that an *ad hoc* correlation of target tree and feeding biology is often premature and incorrect.

### Gut content not matching target tree but potentially pointing to alternative feeding

In four cases (larvae nr. 82, 102, 107, 108, see Table 4) the larvae were fogged down from trees (genus *Paullinia*, Sapindaceae; genus *Annona*, Annonaceae; genus *Apeiba*, Malvaceae), but the cut content was pointing to alternative feeding on mosses (Bryophyta). In the case of the *Lascoria* species (Erebidae; larva nr. 107) such alternative feeding is not excluded as larvae of this species were already observed when grazing on algae in Costa Rica [4]. However, moss-feeding is very unusual in Lepidoptera, and this may also be caused by contamination since these fogging samples (under 80% alcohol) contained some leaves of *Lejeunia* mosses whose DNA may have invaded the larvae through their stigmata or contaminated them on their skin. Further research is needed to estimate the influence of contamination through the sample alcohol. For this purpose larvae should be de-contaminated by bleeching before sequencing. In addition, their gut content could be extracted carefully by cutting the larva longitudinally.

### Inferring potential hostplant relationships (larvae without gut content analysis)

43 larvae with reliable identification to at least genus level, fogged from trees identified to at least genus level give first ‘hints’ on potential host-plants (Table 5). Almost all of them are new records, none of them was found in the ‘Hosts’ database [36] nor in Janzen & Hallwachs [4]. Alternative feeding, however, is not excluded (see notes to larvae nr. 18, 38, 43, 74 and 84 in Table 5), hence all suggested host-plant relationships require confirmation.

**Table 5.** Potential host-plant relationships for 43 larvae identified to at least genus level and identity of the fogged target tree.

| Nr. of larva(e) | Identificarion of larva | Nr. of target tree(s) | Family of target tree | Identity of target tree |
| --- | --- | --- | --- | --- |
|  | **Nymphalidae** |  |  |  |
| 81 | *Memphis acidalia* | 20 | Nyctaginaceae | *Neea* |
| 109 | *Euptychia* n. sp. 5 CP-2006 | 32 | Sapindaceae | *Paullinia* |
|  | **Hesperiidae** |  |  |  |
| 43 | *Panoquina fusina* (1) | 5 | Meliaceae | *Guarea* |
| 115 | *Myscelus epimachia* | 35 | Meliaceae | *Guarea guidonia* |
|  | **Apateodidae** |  |  |  |
| 105 | *Olceclostera* | 32 | Sapindaceae | *Paullinia* |
|  | **Saturniidae** |  |  |  |
| 62 | *Automeris denticulata* | 14 | Moraceae | *Poulsenia/Naucleopsis* |
|  | **Bombycidae** |  |  |  |
| 63 | *Anticla* near *antica*DHJ04 | 14 | Moraceae | *Poulsenia/Naucleopsis* |
| 113 | *Anticla* near *antica*DHJ03 | 34 | Moraceae | *Ficus* |
|  | **Geometridae** |  |  |  |
| 13, 75 | *Hemipterodes divaricata* | 19 | Annonaceae | *Oxandra polyantha* |
| 74 | *Idaea orilochia* | 19 | Annonaceae | *Oxandra polyantha* |
| 38 | *Ergavia* | 1 | Anacardiaceae | *Mangifera indica* |
| 9, 59 | *Thysanopyga apicitruncaria* | 14 | Moraceae | *Poulsenia/Naucleopsis* |
| 58 | *Perissopteryx divisaria* | 14 | Moraceae | *Poulsenia/Naucleopsis* |
| 68 | *Pero incisa* | 17 | Moraceae | *Clarisia biflora* |
| 104 | *Ischnopteris chlorophaearia* | 32 | Sapindaceae | *Paullinia* |
| 116 | *Glena* AH03Pe | 36 | Meliaceae | *Guarea guidonia* |
| 120 | *Physocleora* AH02Pe | 37 | Meliaceae | *Guarea guidonia* |
| 117 | *Semiothisa gambaria* | 36 | Meliaceae | *Guarea guidonia* |
| 123, 130 | *Stegotheca* | 40, 47 | Meliaceae | *Guarea guidonia* |
|  | **Uraniidae** |  |  |  |
| 26 | *Urania leilus* | 27 | Annonaceae | *Oxandra polyantha* |
|  | **Noctuidae** |  |  |  |
| 103 | *Lycaugesia* | 32 | Sapindaceae | *Paullinia* |
|  | **Erebidae Arctiinae** |  |  |  |
| 16 | *Delphyre orientalis* | 20 | Nyctaginaceae | *Neea* |
| 18 | *Prepiella* species 1 (4) | 22 | Malpighiaceae | *Byrsonima coccolobifolia* |
| 66 | *Ernassa* | 15 | Chrysobalanaceae | *Hirtella* |
| 84 | *Clemensia* species 1 (4) | 22 | Malpighiaceae | *Byrsonima coccolobifolia* |
| 99, 100 | *Haemanota nigricollum* | 29 | Malvaceae | *Apeiba* |
| 119 | *Melese* | 37 | Meliaceae | *Guarea* |
|  | **Erebidae other subfamilies** |  |  |  |
| 8 | *Ypsora selenodes* | 14 | Moraceae | *Poulsenia/Naucleopsis* |
| 61 | *Letis magna* | 14 | Moraceae | *Poulsenia/Naucleopsis* |
| 69 | *Lascoria* Poole03 | 17 | Moraceae | *Clarisia biflora* |
| 95 | *Lascoria manes* | 25 | Myristicaceae | *Otoba parvifolia* |
| 71, 73 | *Latebraria amphipyroides* | 19 | Annonaceae | *Oxandra polycarpa* |
| 77 | *Mastixis* | 19 | Annonaceae | *Oxandra polycarpa* |
| 78 | *Feigeria scops* | 20 | Nyctaginaceae | *Neea* |
| 96 | *Clapra* | 27 | Annonaceae | *Oxandra polycarpa* |
| 111 | *Antiblemma sterope* | 33 | Meliaceae | *Trichilia* |
| 114, 126 | *Sosxetra grata* | 35, 41 | Meliaceae | *Guarea* |

(1) potential alternative feeding (members of genus *Panoquina* are known as feeders on monocotyledon plants like Poaceae); (2) potential alternative feeding (members of tribe Idaeini in Europe known as detritus feeders); (3) potential alternative feeding (members of genus *Ergavia* are known as almost exclusively feeding on Polygoniaceae); (4) potential alternative feeding (members of tribe Lithosiini in Europe known as lichenophagous, genus *Clemensia* known as lichenophagous from North America: Host database).

### Target tree confirming previously known host-plants (larvae without gut content analysis)

Among the 87 larvae successfully identified to genus or species and not subjected to gut content analysis, there are at least six cases where the fogged target trees match previously known host-plant relationships: larvae nr. 63 and 113 (Bombycidae, *Anticla* near *antica*) were knocked down from the trees nr. 14 and 34 (Moraceae); larvae nr. 117 (Geometridae; *Semiothisa gambaria*), 115 (Hesperiidae, *Myscelus*) and 144+126 (Erebidae, *Sosxetra grata*) from the trees nr. 35 and 41 (Meliaceae, *Guarea*), all confirming the relationships as previously recorded by Janzen & Hallwachs [4].

### A powerful tool for future synecological research?

Our pilot study has revealed that (1) molecular identification of fogged, neotropical lepidopteran larvae works successfully in general and even down to species level (if already listed in BOLD), that (2) molecular identification of target trees usually works well at least to genus or family level and (3) molecular gut content analysis based on HTS techniques can be used for confirming or rejecting the feeding on the fogged target tree. With further completion of the DNA reference libraries in the future for (1) (Peruvian Lepidoptera; currently 12,746 sequences, 3532 BINs) and (2) (Peruvian plants) a better taxonomic resolution of identification will be achieved, whilst molecular gut content analysis (3) can be improved by de-contamination and/or isolated storage of the fogged larvae.

With that, the herewith presented approach has the potential for unveiling trophic interactions for primary consumers in tropical regions at a very large scale, which can be performed in a fast and cost-effective way considering the steadily dropping costs for DNA barcoding and HTS. The extremely high diversity of 92 species in 119 larvae in our study shows that canopy fogging and molecular analyses may improve synecological knowledge for a broad spectrum of arthropods. The availability of reliable data on trophic interactions is of great importance for forestry, agriculture, biodiversity and ecological research and – last but not least – for conservation purposes. Increasing such knowledge – particularly in megadiverse ecoregions – is an imperative in a world of unprecedented biodiversity losses. In this context, the proposed molecular approach of investigating host-plant relationships constitutes an important research tool, which fits well in the research plan of the recently launched BIOSCAN phase of the international Barcode of Life program ([37]; see also https://ibol.org).

## Acknowledgements

We thank “Moro” Carlos Vásquez Módena (Panguana) and Hamilton Paredes (Museum of Natural History, Lima) for identification of the target trees. We furthermore thank Julio Monzón (Lima / Freiburg) for providing the fogger and the chemicals. Víctor Meyer Ruiz Chota (Yuyapichis) was very helpful in taking the fogging samples, Johanna Brunner, Sarah Augustin and Stefanie Wothe (Munich) helped to take the cambium/sapwood samples. Dr. Andreas Fleischmann (SNSB, Botanische Staatssammlung, Munich) gave advice concerning the botanical aspects of the study. The genetic analyses have received considerable support from Paul D. N. Hebert and the Biodiversity Institute of Ontario (BIO) and the Canadian Centre for DNA Barcoding (CCDB University of Guelph). The data management and analysis system BOLD was provided by Sujeevan Ratnasingham. We thank the Peruvian nature conservation authority and forestry office (Servicio Forestal y de Fauna Silvestre, SERFOR, Ministerio de Agricultura y Riego, MINAGRI) for collection permits (N° 007-2014-SERFOR-DGGSPFFS; N° 0406-2017-SERFOR-DGGSPFFS) and exportation permits (fauna: N° 003281-SERFOR, N° 003320-SERFOR; flora: N° 003284-SERFOR + N° 161-2018-MINAGRI-SERFOR-DGGSPFFS, N° 003333-SERFOR).

## Supporting information

**S1 Table. Morphology-based identification of target trees**. Morphology-based identification of target trees to Peruvian vernacular names (mostly provided by the administrator of the Panguana station, Moro Carlos Vásquez Modena) and attempt to assign scientific family / genus / or species names (partly provided by Hamilton Paredes (“HP”), Museum of Natural History, Lima, based on leaf samples). # = fogging sample from target tree without lepidopteran larva; * = no plant tissue available, so far (hence no molecular confirmation possible): ^1^ = same vernacular name with two different molecular identifications.

**S2 Table. Target Trees: Sequencing success and process identification numbers**. Data from BOLD, with fragment lengths in basepairs (bp). Sanger sequencing of rbcL, trnL-F and psbA genes, based on leaf (l) and cambium+sapwood (c) samples from the target trees.

**S3 Table. Molecular identification of target trees.** Molecular identification of the target trees after Sanger sequencing (rbcL, trnL-F and psbA genes) of leaf (l) and cambium + sapwood (c) samples. Results from blasting on NCBI, BLAST matches (highest percent identity (‘Max ident’) of all query-subject alignments) usually >99.5%, otherwise indicated. Plant species/genera with blast matches sometimes not mentioned when plants are exclusively distributed on other continents. # = fogging sample from target tree without lepidopteran larva in the sample. Anac. = Anacardiaceae; Anno. = Annonaceae; Cann. = Cannabaceae; Chry. = Chrysobalanaceae; Clus. = Clusiaceae; Euph. = Euphorbiaceae; Faba. = Fabaceae; Malv. = Malvaceae; Malp. = Malpighiaceae; Meli. = Meliaceae; Mora. = Moraceae; Myri. = Myristicaceae; Nyct. = Nyctaginaceae; Rubi. = Rubiaceae; Sapi. = Sapindaceae; Sapo. = Sapotaceae; Viol. = Violaceae. ^1^ = same vernacular name with two different molecular identifications; ^2^ = potential sampling error (tissue sampling from neighboring tree or tube flip in the lab process); ^3^ = exclusively Indo-Pacific; ^4^ = exclusively Old World.

